# Development of direction selectivity in the Xenopus tadpole optic tectum and midbrain tegmentum

**DOI:** 10.64898/2026.11.10.765692

**Authors:** Kaiyuan Zheng, Yugarshi Mondal, Bradley P. Roth, Uwemedimo G. Udoh, Ronald L. Calabrese, Kara G. Pratt

## Abstract

Detecting motion direction is a fundamental function performed by direction selective (DS) neurons. DS neurons in the optic tectum (superior colliculus in mammals) enable the tracking of moving objects. DS neurons of the accessory optic system support reflexive eye / head movements that stabilize gaze during self-motion. By developmental stage 48 (10 days postfertilization), Xenopus tadpoles display visually guided behaviors that require both types of DS neurons. Our previous work has shown that these two regions are not only functionally distinct but also follow different developmental programs. Here we characterized and compared how direction selectivity develops in both visual processing centers. Using *in vivo* electrophysiological recordings during moving bar visual stimulation, we measured DS synaptic inputs and spiking outputs in both regions across three key developmental stages of the retinotectal projection. Tegmental neurons were direction selective *ab initio*, and both input and output tuning remained stable across development. Tectal neurons were observed to receive DS inputs early, and their input selectivity also remained stable across development. However, their spiking output grew substantially more DS at the later developmental stage. This was due at least in part to a naturally occurring decrease in intrinsic excitability which preferentially dampened overall spike rates, but especially in response to the non-preferred direction of motion, thereby increasing direction selectivity. These results reveal distinct developmental trajectories in parallel visual pathways. Tegmental direction selectivity is established early and stays stable, whereas tectal output tuning sharpens during development through a decrease in intrinsic excitability.

## Introduction

The ability to perceive direction of motion is a fundamental feature of any visual system. Direction of motion is computed by direction selective (DS) neurons, neurons that are most strongly activated by a specific direction of motion. DS neurons are known to reside across many visual processing centers. Two main components of the vertebrate visual system, the optic tectum (superior colliculus in mammals) and the nuclei of the accessory optic system (AOS), are known to contain DS neurons (Rasmussen and Yonehara, 2020). The DS neurons in these two midbrain visual processing centers carry out distinct functions. In the optic tectum / superior colliculus, DS neurons contribute to identifying and tracking objects in space. DS neurons of the AOS contribute to the sensory side of the optokinetic reflex – self motion-triggered compensatory eye movements that stabilize the visual scene on the retina. This stabilization prevents blur and allows other higher order components of the visual system to function properly. Studies in mouse have established that DS neurons in these two regions inherit direction selectivity directly from different types of DS retinal ganglion cells (RGCs; Elstrott and Feller, 2009; Sun et al., 2015; Shi et al., 2017; Harris and Dunn, 2023).

Here, we characterize the development of direction selectivity in the *Xenopus laevis* optic tectum and basal optic nucleus (BON), a group of neurons residing in the ventral midbrain tegmentum which, together with the presynaptic retinal ganglion cell input, constitute the amphibian AOS (Figure 1A; Sakaguchi et al., 1985; Udoh et al., 2025). The Xenopus tadpole is especially well-suited for this study because *in vivo* whole cell recordings can readily be carried out from neurons of both the optic tectum and midbrain tegmentum, across well described developmental stages that span from developmental stage 42 (5-6 days postfertilization; dpf) when these circuits are just beginning to wire up and tadpoles are mostly immobile, through developmental stage 48 (10-18 dpf), the time by which the circuits are well formed and tadpoles are displaying exploratory swimming and several different innate visually guided behaviors (Dong et al., 2009; Hunt et al., 2020; Bruno et al., 2022; Udoh et al., 2025; Barrus et al., 2026; Reviewed by Hayes et al., 2026).

**Figure 1:**
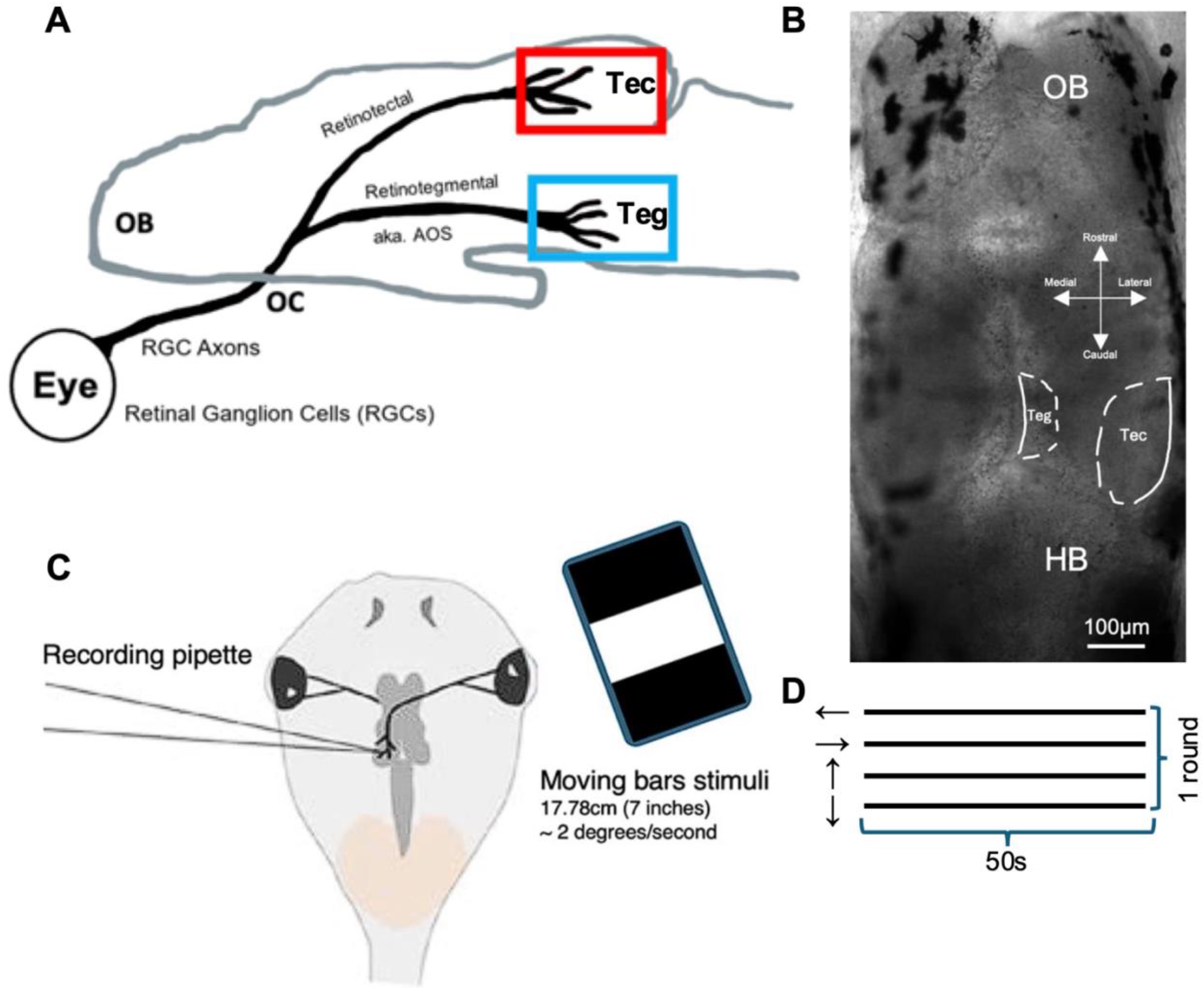
I*n vivo* recording experimental set up. **(A)** Sagittal diagram showing the two distinct projections of RGC axons coming from the back of the eye, innervating to two of its midbrain targets, the optic tectum and the tegmentum. **(B)** Microscopy image of a filleted *in vivo* tadpole brain taken at 5x amplification showing the real-life relative position of the optic tectum and the tegmentum. **(C)** Schematic showing the *in vivo* recording set-up. The recording pipette is set to record from either the midbrain optic tectum or the tegmentum contralateral to the eye that is projected the moving bars of stimuli. The moving bars are 7 inches in width, moving at 2 angular degrees per second speed. The screen is positioned perpendicularly to the eye contralateral to the midbrain being recorded from, 50cm away.**(D)** Schematic showing the visual stimuli protocol for measuring direction selectivity. One 50 seconds trial for each of the four orthogonal directions. One round contains four trials of each direction. For synaptic input recordings, 3 rounds of recordings were carried out; for spike output recordings, 2 rounds of recordings were carried out. RGCs: retinal ganglion cells; OB: olfactory bulb; OC: optic chiasm; Tec: optic tectum; Teg: tegmentum; AOS: accessory optic system; μm: micrometer; HB: Hind brain; cm: centimeter; s: seconds; ←: temporal to nasal; →: nasal to temporal; ↑: upward; ↓: downward.

Direction selectivity of synaptic input was measured by recording excitatory post synaptic currents (EPSCs) in response to a bar stimulus moving in 4 orthogonal directions. Direction selectivity of action potential spike output was measured from a different set of tectal and tegmental neurons by recording spikes in loose cell attached mode in response to the same moving bar stimulus.

Overall, we found a consistent population of tegmental neurons that received DS input and that displayed DS output across all three stages of development. Furthermore, the strength of direction selectivity, also referred to as direction tuning, was also stable across development, suggesting functional direction selectivity in the AOS as soon as the RGC axonal inputs form functional synapses with postsynaptic tegmental neurons. Development of direction selectivity was observed to be considerably more dynamic in the optic tectum. The percentage of tectal neurons that received DS input varied dramatically across development. We also observed that while the direction selectivity of synaptic input received by tectal neurons appeared broadly tuned across development, DS spike output sharpened significantly between developmental stage 45 and 48. Combined, our data suggest that the sharpening of direction tuning at the later stage was manifested through a naturally occurring increase in tectal neuron spike threshold at development stage 48.

## Methods

### Animal care and *in vivo* preparation

All experimental protocols involving animals were approved by the Institutional Animal Care and Use Committee (IACUC) of the University of Wyoming. Wild-type *X. laevis* embryos obtained by in-house mating were reared in 10% Steinberg’s solution at 22°C on a 12-hour light/dark cycle. Tadpoles were staged according to the method of Nieuwkoop and Faber (1994). Recordings were carried out across three key developmental stages: NF stages 42 (∼5-6 days postfertilization), 45 (∼7-8 days postfertilization), and 48 (∼10-18 days postfertilization). Tadpole sex is indeterminate at these early stages of development.

Tadpoles were anesthetized with 0.01% tricaine methanesulfonate (MS-222) and transferred to a recording dish that contains HEPES-buffered extracellular saline (composition in mM: 115 NaCl, 4 KCl, 3 CaCl_2_, 3 MgCl_2_, 5 HEPES, 0.01 glycine and 10 glucose; pH 7.25; osmolarity 255mOsm) and containing the paralytic tubocurarine (0.01mM) to prevent muscle fiber contractions during recording. The tadpole is stabilized by insect pins onto a piece of submerged Sylgard silicone elastomer in the recording chamber. To access neurons for recording, the skin above the brain was peeled away and the entire brain was filleted along the dorsal midline.

#### Imaging

The image of *In vivo* filleted brain of a tadpole was acquired using an upright Zeiss laser scanning confocal microscope (Figure 1B; LSM 700, Carl Zeiss AG, Germany). The image was further cropped using ImageJ (NIH, Bethesda, MD, US) software version 1.53k.

### Moving stimuli

Alternating black and white slow-moving bars of stimuli moving in four orthogonal directions: temporal (T) to nasal (N), nasal to temporal, upward, downward were projected onto one eye. The temporal to nasal and nasal to temporal directions are the horizontal orientation. The upward and downward directions are the vertical orientation. The moving bars stimuli were displayed via a wired computer screen (15.6”, 16:9, 1080p 60Hz, Innocn, China) connected to a computer. The screen is perpendicularly positioned 50cm away from the eye contralateral to the recording side of the brain. The moving bars of stimuli are 7” wide and move at 2° per second across the retina (Figure 1C). DS neurons in the vertebrate AOS are known to preferentially respond to slow, wide field motion (Burns and Wallman, 1981; McKenna and Wallman, 1985; Soodak and Simpson, 1988). Accordingly, we used slow-moving, relatively wide bars, which additionally minimized the risk of any potential stimulus-induced plasticity. Moving bars in each of the four orthogonal directions was presented to the tadpole for 50 seconds each per trial (Figure 1D). Within this 50 second duration, 10 borders of bars pass by the entire screen. Depending on the recording protocol, either two or three rounds of trials were run, further details explained below.

### *In vivo* electrophysiological recordings

For our recordings, we defined the tegmentum as the most ventral part of the midbrain opposite the widest part of the tectum. In our previous work be characterized the formation of this accessory optic system, including mapping the position in the midbrain tegmentum, the position of the BON neurons, which we define as the neurons in the tegmentum that receive direct RGC input (Udoh et al., 2025). Hence, for this study, we restricted our recordings of tegmental neurons within the region of the tegmentum that we established as most likely being BON neurons (Figure 1B). We restricted our recordings of tectal neurons to those residing in the middle third of the optic tectum (OT) along the rostral-caudal (R-C) axis. This controls for variation due to the developmental gradient that is known to exist across the optic tectum’s R-C axis (Wu et al., 1996; Hamodi et al., 2016; Udoh et al., 2025). For each preparation, either the tegmentum or the optic tectum is recorded from. For accessing neurons in either the tegmentum or the optic tectum, the skin overlying the tadpole brain was removed and the brain filleted along the midline to expose the ventral surface of the midbrain, then the ventricular membrane above selected brain region was removed by using a broken borosilicate glass micropipette. Recordings were carried out at room temperature (20-22°C) with borosilicate glass micropipettes with resistance in the range of 7-14MΩ, containing potassium gluconate internal saline (composition in mM: 100 potassium gluconate, 8 KCl, 5 NaCl, 1.5 MgCl_2_, 20 HEPES, 10 EGTA, 2 ATP, and 0.3 GTP; pH 7.2; osmolarity 255mOsm).

All *in vivo* electrophysiological recordings were carried out with Axon Instruments MultiClamp 700B microelectrode amplifier and digitized at 10kHz using Digidata 1550B digitizer (Molecular Devices, San Jose, CA, US), paired with the pClamp software version 11.2 and Multiclamp 700B Commander software version 2.2.1 (Molecular Devices, San Jose, CA, US) on a Dell PC. Tegmental and tectal neurons were visualized using a Zeiss AXIO Examiner.A1 light microscope with a 63x/1.0 water immersion objective that is connected to a Hamamatsu Complementary Metal-Oxide-Semiconductor (CMOS) digital camera (ORCA-spark, Hamamatsu Photonics K.K., Japan).

#### Measuring direction selectivity of synaptic input

To measure direction selectivity of synaptic input, whole cell electrophysiological recordings were carried out *in vivo* in voltage-clamp mode at -45mV, the reversal potential of gamma-aminobutyric acid (GABA) current activities, to exclusively collect only excitatory current activities. Typically, 4 of either tegmental or tectal neurons per experimental preparation were recorded from a whole cell configuration. Voltage-clamp recordings were carried out with 2 x 50 seconds of stationary, colored-image baseline and following 3 rounds of 50 seconds trials of moving bars in each of the four orthogonal directions.

#### Measuring direction selectivity of output

To measure direction selectivity of action potential (AP) spiking output, *in vivo* loose cell-attached recordings were carried out to quantify output spike counts of neurons due to the moving bars of stimuli. To achieve a loose cell-attached configuration, we acquired a 70∼100MΩ loose seal onto the neuronal membrane, and recordings were carried out in current-clamp mode. Typically, 6 of either tegmental or tectal neurons per experimental preparation were recorded from a loose cell-attached configuration. Loose cell-attached recordings were carried out with 1 x 50 seconds of stationary, colored-image baseline and following 2 rounds of 50 seconds trials of moving bars in each of the four orthogonal directions.

To quantify peak Na^+^ and K^+^ currents, neurons were held at -60mV in an *in vivo* whole-cell voltage-clamp configuration, stepped to increasingly more depolarized potentials, from -60mV to 30mV, for a duration of 250ms each, and the resulting voltage-dependent Na^+^ and K^+^ currents were recorded. Leak current was subtracted by pClamp software in real time.

Resting membrane potential (RMP) was determined by switching out of voltage-clamp mode to “*I* = 0” mode and waiting an average of 5 seconds to get a stable RMP. The RMP has not been adjusted for a -12mV liquid junction potential. Cells with > 50MΩ access (series) resistances and/or > -50pA holding current were not included in the dataset.

To measure AP firing thresholds, whole cell current-clamp ramp tests were carried out to measure the AP firing thresholds of stage 45 and stage 48 tectal neurons. During ramp tests, neurons were put in current-clamp mode and injected currents to keep them initially held at around -40∼-55 mV. Within the ramp test protocol, tectal neurons were injected with 10pA of currents after a 100ms delay, over 500ms duration, for 10 ramps with increasing currents in total.

### Pharmacological interventions

All pharmacological agents used in this study were prepared as stock solutions, aliquoted, and stored at -20°C until use. Tadpoles are exposed to 2-amino-5-phosphonovaleric acid (APV, 100μM, Tocris, UK, CAT#0106) in their rearing solution at developmental stage 45. The concentrations used were the highest concentrations that did not elicit any noticeable effects on normal swimming patterns and behavior. APV is a highly selective and potent competitive NMDA receptor antagonist, and it is known to block any activity-dependent synaptic refinement (Udoh et al., 2025). Electrophysiological recordings of action potential spikes caused by moving bars of stimuli are carried out in loose cell-attached mode at stage 48 *in vivo*. Batch matched controls were also recorded.

Tadpoles were exposed to picrotoxin (100µM, Acros Organics, Belgium, CAT#131210050) in their rearing solution at stage 42 (Deeg and Aizenman, 2011), or, applied to the external recording solution acutely (Udoh et al., 2024). Picrotoxin is a drug that blocks GABAa receptors, thus by rearing tadpoles with it would abolish inhibition dependent tuning effects. Electrophysiological recordings of action potential spikes caused by moving bars of stimuli are carried out in loose cell-attached mode at stage 48 in vivo. Batch-matched controls were also recorded.

### Statistical analysis

All the electrophysiological recording traces were analyzed off-line with AxoGraph (AxoGraph Scientific, Australia) software version 1.8.0 and graphed using Igor Pro (WaveMetrics Inc., Portland, OR, US) software version 9.05. Synaptic events were collected and quantified using a variable amplitude template (Clements and Bekkers, 1997). We used the synaptic drive index (SDI) to estimate the strength of excitatory input for each direction in each neuron by multiplying the average excitatory postsynaptic current (EPSC) amplitude by the average EPSC frequency (Pratt and Aizenman, 2007):

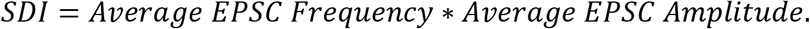

The average EPSC frequency for each neuron was calculated by dividing the total number of synaptic events recorded across three 50 s trials by the total duration (150 s) to obtain the event rate in Hertz (events/s). The average EPSC amplitude for each neuron was determined by averaging the event amplitudes across the three 50 s trials presented in the same direction (Figure 1D). The DSI equation used to calculate the strength of a neurons’ direction selectivity for the two directions within the same vertical or horizontal orientation is (Mazurek et al., 2014)

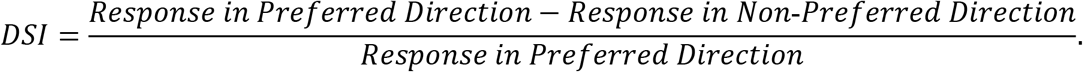

For synaptic input, the average SDIs of the two directions from the same orientation were used to calculate the input DSI; For spike output, the absolute number of spikes of the two directions from the same orientation were used to calculate the output DSI.

Neurons with a DSI ≥ 0.2 were considered direction selective. For cells that were direction selective along both the horizontal and vertical orientations, both of their DSI values were included. Because the leading and trailing borders of the moving bars crossed the screen ten times during each 50 s trial, we applied an additional inclusion criterion: a neuron was retained only if it produced, on average, at least ten events (ten synaptic events or ten spikes per 50 s trial; 0.2 Hz) in at least one of the four cardinal directions. We refer to this throughout the Results as the "event count criterion."

Descriptive and inferential statistical analyses were carried out using GraphPad Prism (GraphPad Software, Boston, MA, US) software version 11.0.2 and Microsoft Excel (Microsoft, Redmond, WA, US) software version 16.105.3. Schematic graphs were conceived and graphed in Microsoft PowerPoint software version 16.111.3 and cropped in Preview (Apple, Cupertino, CA, US) software version 11.0. All experimental graphs and figures were conceived and graphed using Igor Pro. All descriptive statistics are reported as mean ± standard error of the mean (SEM).

Differences in means between groups of measurements (e.g., DSI, SDI, or spike count) were assessed using two-tailed, unpaired Welch’s t-tests when neither group showed a significant departure from normality under the Shapiro–Wilk test. Otherwise, a two-tailed Mann–Whitney U test was used to assess differences between the distributions of measurements in the two groups. Statistical differences with a \**p* value <0.05 were considered statistically significant. Exact *p* values are reported where available; *p* values below the numerical reporting limit of GraphPad Prism are reported as \*\*\*\**p*<0.0001. Unless indicated otherwise, summary data is presented using a dot plot.

## Results

Direction selectivity of synaptic input was measured by recording responses of individual neurons to slowly drifting bars of light, moving in four orthogonal directions (temporal to nasal, nasal to temporal, upward, and downward). Resulting excitatory postsynaptic currents (EPSCs) were isolated by recording neurons voltage-clamped at -45 mV, the reversal potential for GABA-mediated inhibitory synaptic currents. The amount of excitatory synaptic input elicited by each direction of motion was quantified by calculating the SDI, the product of EPSC frequency and the average EPSC amplitude (which reflects synaptic strength) evoked by each direction. From a different set of tegmental and tectal neurons, direction selectivity of action potential (“spike”) output was measured using the same stimuli and recording in loose cell attached configuration.

Direction selectivity of both synaptic input and spike output was quantified by calculating the direction selectivity index (DSI). Neurons with a DSI ≥ 0.2 were classified as direction selective (DS). DSI values of synaptic input and action potential output were calculated for tegmental and tectal regions across three key stages of retinotectal circuit formation: developmental stage 42 (approximately 5-6 days postfertilization, dpf), when the retinal ganglion cell (RGC) axons have just reached the midbrain and have begun forming nascent synapses, stage 45 (approximately 7-8 dpf) when the circuit is moving through a rapid phase of refinement, and stage 48 (approximately 10-18 dpf), a stage at which the retinotectal projection has completed one phase of refinement, resulting in improved visual acuity. (Our previous work revealed that while the retinotectal projection is moving through these dynamic phases of development, the retinotegmental projection remains stable from its earliest stage of formation, stage 42; Udoh et al., 2025). The following is a report on our findings on the development of direction selectivity of synaptic input and spike output, first of tegmental and then tectal neurons.

### A subset of tegmental neurons was observed to receive DS input from the earliest stage of retinotegmental circuit formation

Figure 2A shows an example of a recording from a tegmental neuron which is receiving DS input in the horizontal orientation. Based on the 0.2 DSI threshold, we identified tegmental neurons that received DS synaptic input from the earliest stage of retinotegmental circuit formation, developmental stage 42 (Figure 2B). Figure 2B shows DSI values for each neuron recorded, in both the vertical and horizontal orientations, at the three developmental stages. The percentage of tegmental neurons that received DS synaptic input remained relatively stable across development (at stage 42, 33% of recorded neurons that met the event count criterion were classified as DS; at stage 45, 38% were DS; and at stage 48, 29%; Table 1). To further characterize direction selective synaptic input, we plotted the synaptic drive indexes (the estimate of total synaptic input which is used to calculate the DSI) across the three stages of development (Figure 2C). Among tegmental neurons receiving DS input, we found no statistically significant differences in average DSI values across the three developmental stages (Figure 2D, Table 1), suggesting that the strength of DS synaptic input received by tegmental neurons is remaining stable across development. The unchanging tuning of DS synaptic input received by tegmental neurons across development is consistent with our previous finding that the retinotegmental synapse is relatively hardwired (Udoh et al., 2025). Also, that direction selectivity is observed *ab initio*, as soon as the retinotegmental projection forms, suggests that the source of DS input received by tegmental neurons is likely directly from DS retinal ganglion cells, as has been reported for DS input to the nuclei of the midbrain AOS (Giolli et al., 2006; Krause et al., 2014; Sun et al., 2015; Bastakov et al., 2015).

**Figure 2:**
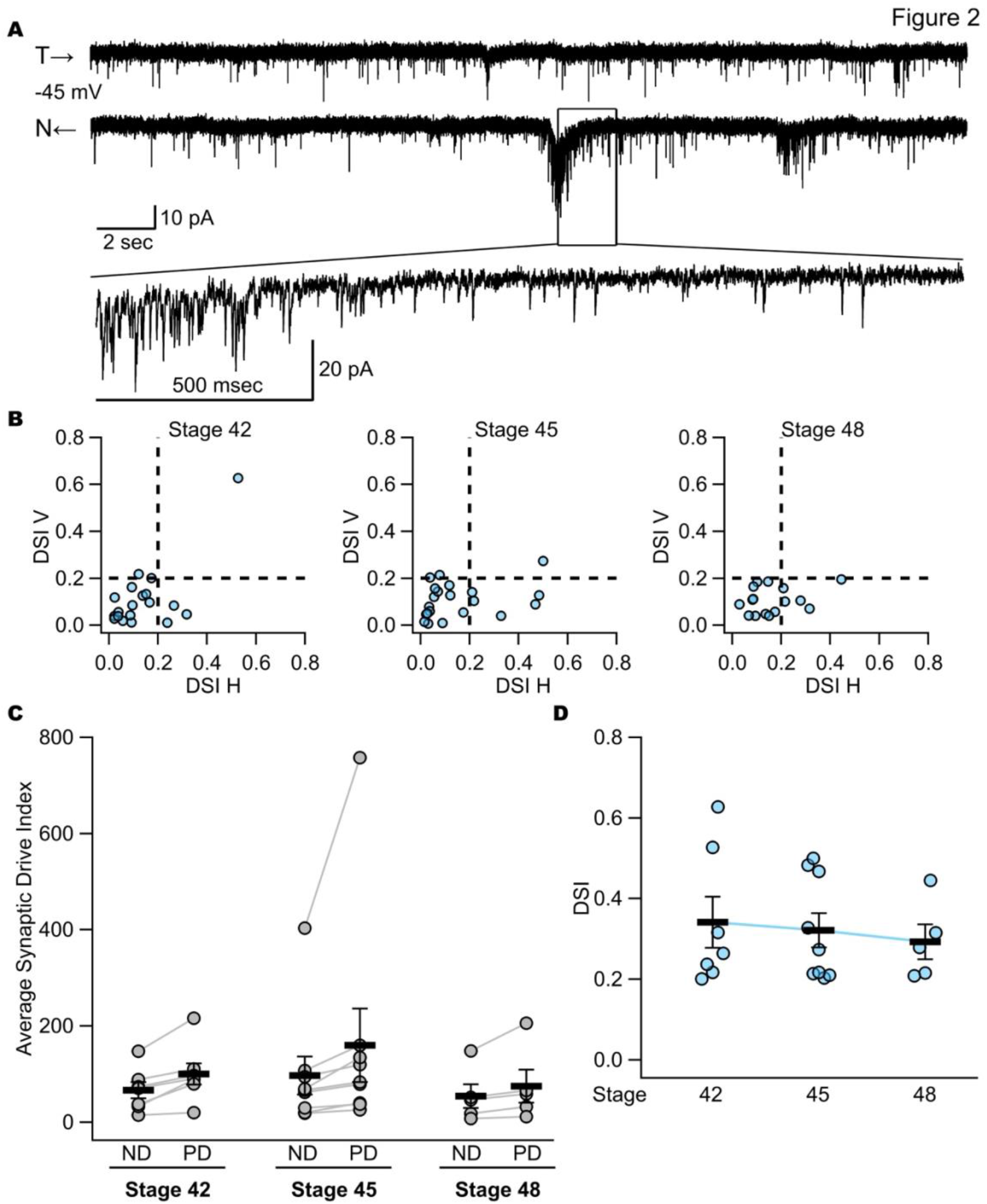
The synaptic input direction selectivity of tegmental neurons does not vary much across developmental stages. **(A)** 30 second chunk with a zoomed in 2 second snippet of a sample recording trace of a voltage-clamp recording recorded at -45mV, from a stage 42 direction selective tegmental neuron, preferring the temporal to nasal direction. **(B)** Dot plots representing the relative strength of direction tuning, the DSI, of synaptic inputs that stage 42, 45, and 48 midbrain tegmental neurons received when moving bars are projected to the contralateral eye. Each dot represents a single neuron that has met the events count criterion, displaying its DSI value for both vertical and horizontal orientations. The dash line represents the deliberate 0.2 DSI cut off. The proportion of DS tegmental neurons remained stable across development: stage 42: 33%, qualified cell n=18, 6 were DS; stage 45: 38%, qualified cell n=21, 8 were DS; stage 48: 29%, qualified cell n=17, 5 were DS. **(C)** Dot plot showing the average SDI of each tegmental neuron, paired in non-preferred direction versus preferred direction across three developmental stages. For neurons displaying direction selectivity in both orientations, both pairs of their SDI values are included in the calculations. The bars represent the SDI mean ± SEM. At stage 42, the average SDI for ND is 66.70±16.77; for PD is 100.51±22.19. At stage 45, the average SDI for ND is 96.98±39.72; for PD is 159.97±76.35. At stage 48, the average SDI for ND is 54.70±24.90; for PD is 75.15±34.12. **(D)** Categorical trend plot depicting the overall trend of input DSI for only direction selective tegmental neurons across three key developmental stages. For neurons displaying direction selectivity in both orientations, both of their DSI values are included in the calculations. A prime example of this is at stage 42 where we have one neuron that is strongly direction selective in both orientations, it shows in 2B, and since both orientations are direction selective for this cell, it shows two points in 2D. At stage 42, the average input DSI of tegmental neurons at stage 42 is 0.34±0.06, n=7; at stage 45 is 0.32±0.04, n=9; and at stage 48 is 0.29±0.04, n=5. For between group comparisons, stage 42 vs. stage 45 tegmental DSI is *p*=0.8371, n.s., Mann-Whitney U test; stage 45 vs. stage 48 tegmental DSI is *p*=0.7972, n.s., Mann-Whitney U test. The bars represent the mean ± SEM. T→: nasal to temporal; N←: temporal to nasal; mV: millivolt; sec: seconds; msec: millisecond; pA: picoampere; DSI: direction selective index; V: vertical; H: horizontal; ND: non-preferred direction; PD: preferred direction.

**Table 1:**
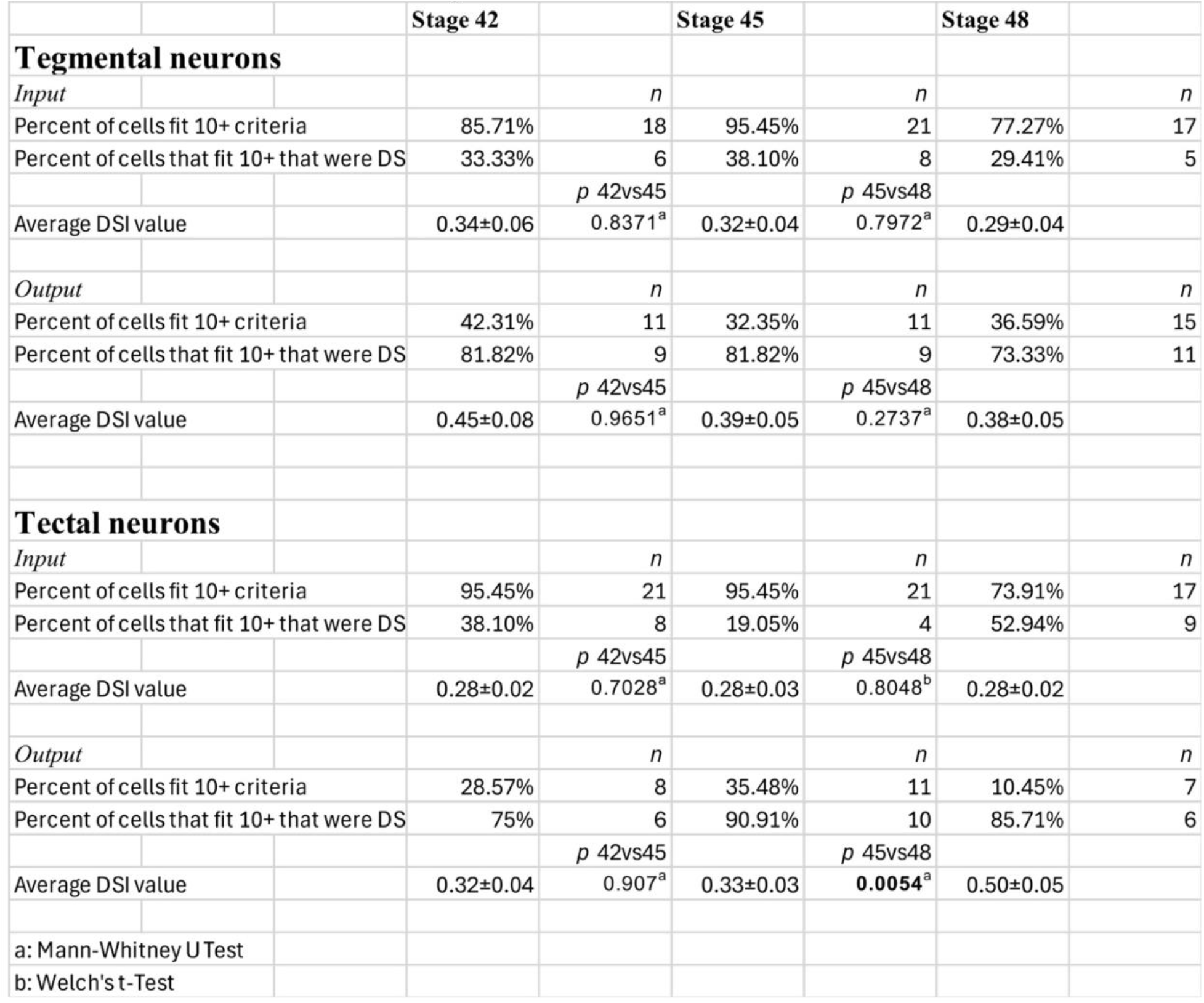
Direction selective statistics. a: Mann-Whitney U Test; b: two-tailed unpaired Welch’s *t*-test, data fit Gaussian distribution, thus unpaired Welch’s *t*-test was used. Abbreviations: DS: direction selective; DSI: direction selective index; vs: versus; n: number of cells.

|  |  |  | **Stage 42** |  | **Stage 45** |  | **Stage 48** |  |
|---|---|---|---|---|---|---|---|---|
| **Tegmental neurons** |  |  |  |  |  |  |  |  |
| *Input* |  |  |  | *n* |  | *n* |  | *n* |
| Percent of cells fit 10+ criteria |  |  | 85.71% | 18 | 95.45% | 21 | 77.27% | 17 |
| Percent of cells that fit 10+ that were DS |  |  | 33.33% | 6 | 38.10% | 8 | 29.41% | 5 |
|  |  |  |  | *p* 42vs45 |  | *p* 45vs48 |  |  |
| Average DSI value |  |  | 0.34±0.06 | 0.8371<sup>a</sup> | 0.32±0.04 | 0.7972<sup>a</sup> | 0.29±0.04 |  |
| *Output* |  |  |  | *n* |  | *n* |  | *n* |
| Percent of cells fit 10+ criteria |  |  | 42.31% | 11 | 32.35% | 11 | 36.59% | 15 |
| Percent of cells that fit 10+ that were DS |  |  | 81.82% | 9 | 81.82% | 9 | 73.33% | 11 |
|  |  |  |  | *p* 42vs45 |  | *p* 45vs48 |  |  |
| Average DSI value |  |  | 0.45±0.08 | 0.9651<sup>a</sup> | 0.39±0.05 | 0.2737<sup>a</sup> | 0.38±0.05 |  |
| **Tectal neurons** |  |  |  |  |  |  |  |  |
| *Input* |  |  |  | *n* |  | *n* |  | *n* |
| Percent of cells fit 10+ criteria |  |  | 95.45% | 21 | 95.45% | 21 | 73.91% | 17 |
| Percent of cells that fit 10+ that were DS |  |  | 38.10% | 8 | 19.05% | 4 | 52.94% | 9 |
|  |  |  |  | *p* 42vs45 |  | *p* 45vs48 |  |  |
| Average DSI value |  |  | 0.28±0.02 | 0.7028<sup>a</sup> | 0.28±0.03 | 0.8048<sup>b</sup> | 0.28±0.02 |  |
| *Output* |  |  |  | *n* |  | *n* |  | *n* |
| Percent of cells fit 10+ criteria |  |  | 28.57% | 8 | 35.48% | 11 | 10.45% | 7 |
| Percent of cells that fit 10+ that were DS |  |  | 75% | 6 | 90.91% | 10 | 85.71% | 6 |
|  |  |  |  | *p* 42vs45 |  | *p* 45vs48 |  |  |
| Average DSI value |  |  | 0.32±0.04 | 0.907<sup>a</sup> | 0.33±0.03 | **0.0054**<sup>a</sup> | 0.50±0.05 |  |
| a: Mann-Whitney U Test |  |  |  |  |  |  |  |  |
| b: Welch's t-Test |  |  |  |  |  |  |  |  |

### Direction selective action potential (spike) output displayed by tegmental neurons mirrors their direction selective synaptic input

Because direction selective input is only functionally meaningful if it is converted into downstream signaling and potential motor outputs, we next asked whether these synaptic responses are reflected in neuronal spiking output. We measured output direction selectivity by recording action potential responses to the same four directions of moving bars using loose cell-attached recordings. Figure 3A shows an example of a tegmental neuron that was direction selective in the vertical orientation. Consistent with the synaptic input, tegmental neurons displaying DS output were present from the earliest stage of circuit formation (examples shown in Figure 4A-C). Among the recorded tegmental neurons that met the event count criterion, 82% were DS at stage 42; 82% at stage 45; and 73% at stage 48 (Figure 3B, Table 1). Figure 3C shows the average spike count per 50 second trial elicited by the preferred and non-preferred direction of motion per 50 second trial. The difference between the average spike counts per trial evoked by the preferred and non-preferred direction of motion was similar across stages (Figure 3C), and this led to consistent DSI values (Figure 3D). Among tegmental neurons displaying DS output, we found no statistically significant differences in average DSI values across the three developmental stages (Figure 3D, Table 1), suggesting that the strength of DS output displayed by tegmental neurons is remaining stable across development. Comparing the average DSI value of tegmental input and output across development shows that they are closely matched, however the spike output DSI had a trend of being consistently slightly higher compared to the corresponding synaptic input DSI (Figures 2D and 3D, Table 1). This may be due to the filtering effect of the spike threshold, which is known to boost direction tuning of spike output (Jagadeesh et al., 1997; Priebe and Ferster, 2008). Overall, for tegmental neurons, the DS synaptic input coming in is broadly tuned, and the resulting output is broadly tuned, too. This concludes our report on the development of direction selectivity in the midbrain tegmentum.

**Figure 3:**
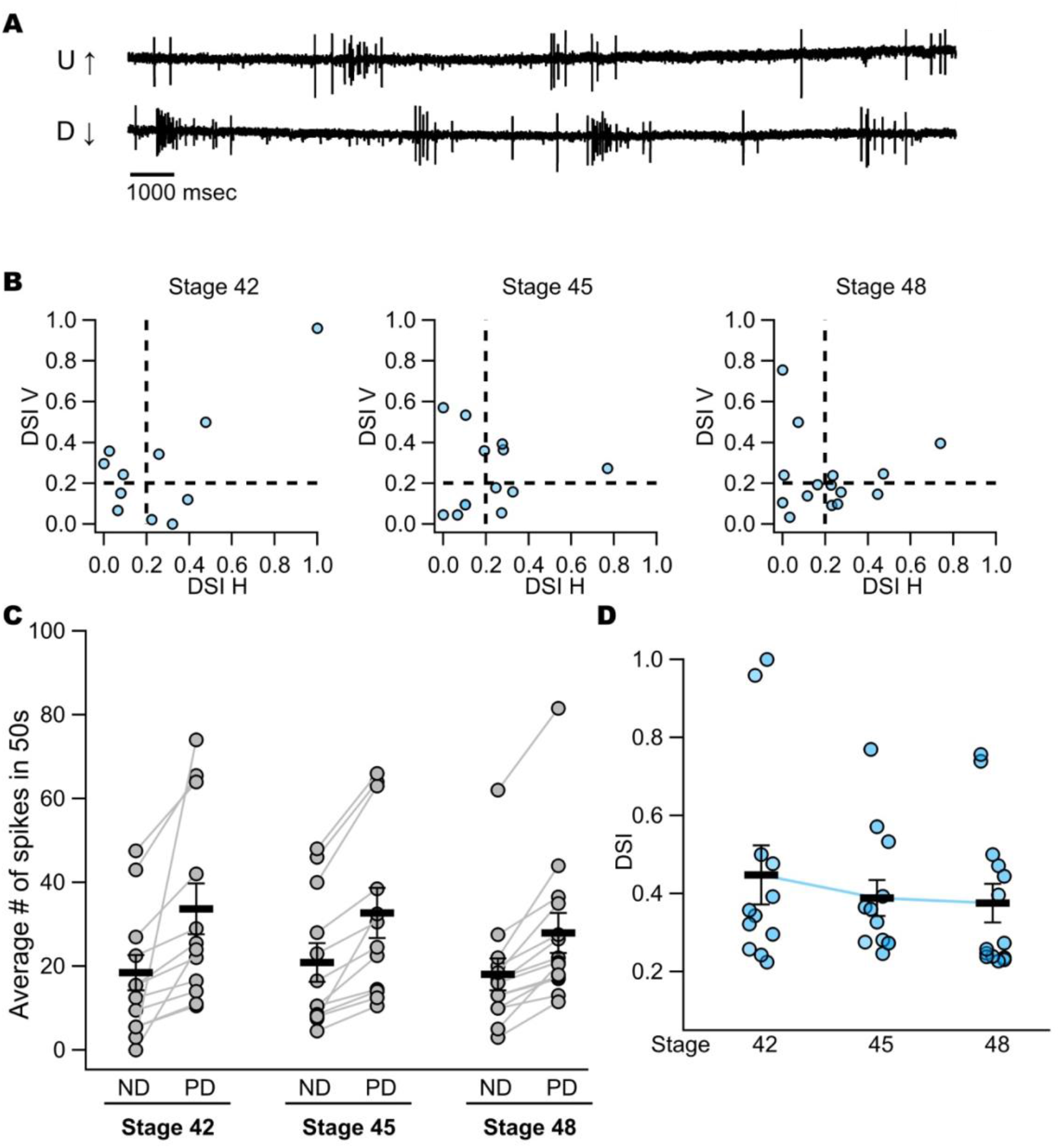
The output direction selectivity of tegmental neurons exhibits developmental program similar to its input across stages. **(A)** 20 second snippet sample trace of a loose cell-attached recording, from a stage 48 direction selective tegmental neuron, preferring the upward to downward direction. **(B)** Dot plots representing the relative strength of direction tuning, the DSI, of output action potentials that stage 42, 45, and 48 midbrain tegmental neurons fired when moving bars are projected to the contralateral eye. Each dot represents a single neuron that has met the event count criterion, displaying its DSI value for both vertical and horizontal orientations. The dash line represents the deliberate 0.2 DSI cut off. For neurons in the tegmentum, about 82% of tegmental neurons are DS at stage 42, qualified cell n=11, 9 were DS; about 82% of tegmental neurons are DS at stage 45, qualified cell n=11, 9 were DS; about 73% of tegmental neurons are DS at stage 48, qualified cell n=15, 11 were DS. **(C)** Dot plot showing average number of spikes in 50 seconds of each tegmental neuron, paired in non-preferred direction versus preferred direction across three developmental stages. For neurons displaying direction selectivity in both orientations, both pairs of their spike count per 50 seconds trials are included in the calculations. The bars represent mean ± SEM of the average number of spikes in each 50 second trial. At stage 42, the average spiking for ND is 18.50±4.24; for PD is 33.65±6.05. At stage 45, the average spiking for ND is 20.92±4.60; for PD is 32.75±6.03. At stage 48, the average spiking for ND is 18.07±3.80; for PD is 27.96±4.81. **(D)** Categorical trend plot depicting the overall trend of output DSI for only direction selective tegmental neurons across three key developmental stages. For neurons displaying direction selectivity in both orientations, both of their DSI values are included in the calculations. The average DSI of tegmental neurons at stage 42 is 0.45±0.08, n=12; at stage 45 is 0.39±0.05, n=12; and at stage 48 is 0.38±0.05, n=14. For between group comparisons, Mann-Whitney U test of stage 42 vs. stage 45 tegmental DSI is *p*=0.9651, Mann-Whitney U test of stage 45 vs. stage 48 tegmental DSI is *p*=0.2737. The bars represent the mean ± SEM. U↑: upward; D↓: downward; msec: millisecond; DSI: direction selective index; V: vertical; H: horizontal; s: seconds; ND: non-preferred direction; PD: preferred direction.

**Figure 4:**
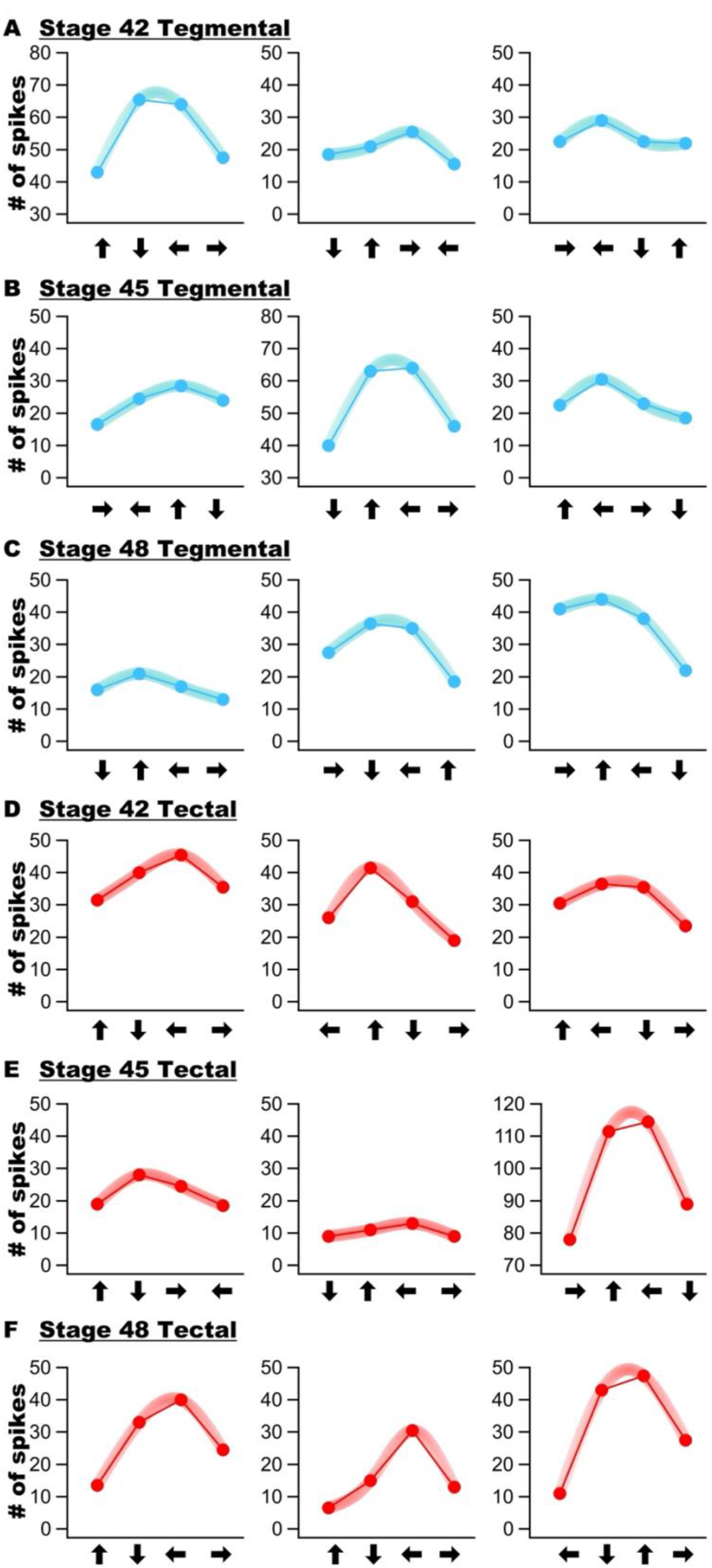
The interpolated direction tuning curves of stage 48 tegmental neurons, and stage 48 tectal neurons show distinct direction tuning sharpness. **(A)** Example tuning curves interpolated from three randomly chosen stage 42 direction selective tegmental neurons. **(B)** Example tuning curves interpolated from three randomly chosen stage 45 direction selective tegmental neurons. **(C)** Example tuning curves interpolated from three randomly chosen stage 48 direction selective tegmental neurons. **(D)** Example tuning curves interpolated from three randomly chosen stage 42 direction selective tectal neurons. **(E)** Example tuning curves interpolated from three randomly chosen stage 45 direction selective tectal neurons. **(F)** Example tuning curves interpolated from three randomly chosen stage 48 direction selective tectal neurons. Each of the arrows represents respective directions of the moving bars: ←: temporal to nasal; →: nasal to temporal; ↑: upward; ↓: downward.

### A subset of tectal neurons was observed to receive DS input across all three developmental stages

We found that a subset of tectal neurons also received DS synaptic input by developmental stage 42, the earliest stage of retinotectal circuit formation (Figure 5A, example trace of a stage 42 tectal DS neuron, receiving direction selective inputs in the vertical orientation). But unlike the consistent percentage of tegmental neurons that were observed to receive DS input across development, the percentage of tectal neurons whose motion-evoked synaptic input qualified as DS varied markedly across development: we observed a statistically significant decline in the percentage of tectal neurons that received DS input between stages 42 and 45, followed by an even more profound increase between developmental stage 45 and 48 (at stage 42, 38% of recorded neurons that met the event count criterion were classified as DS; at stage 45, 19% were DS; and at stage 48, 53%; Table 1, Figure 5B). To further characterize direction selective synaptic input as we did with tegmental neuron analysis, we plotted the synaptic drive index values for each of the DS inputs across the three stages of development (Figure 5C). Among the tectal neurons that received DS input, for their average amount of synaptic drive elicited by both the preferred and non-preferred direction, we observed a statistically significant increase between developmental stages 42 and 45 (\**p*=0.0231, n=26 for stage 42, n=10 for stage 45), and then a statistically significant decrease between stages 45 and 48 (\*\*\**p*=0.0005, n=10 for stage 45, n=24 for stage 48, Figure 5C). While we observed variations in overall amounts of synaptic input across development, the relative difference between amount of synaptic input evoked by the preferred and non-preferred directions was similar, giving rise to statistically indistinguishable average DSI values (Figure 5D). This indicates that the direction selectivity of excitatory synaptic input does not become more strongly tuned across development. In summary, we observed a marked increase in the percentage of tectal neurons receiving DS input between stages 45 and 48 (Table 1; Figure 5B). At the same time, the degree of direction tuning stays stable across the three developmental stages (Figure 5D) in the face of major changes in absolute amounts of synaptic input evoked by the moving bar stimulus (Figure 5C).

**Figure 5:**
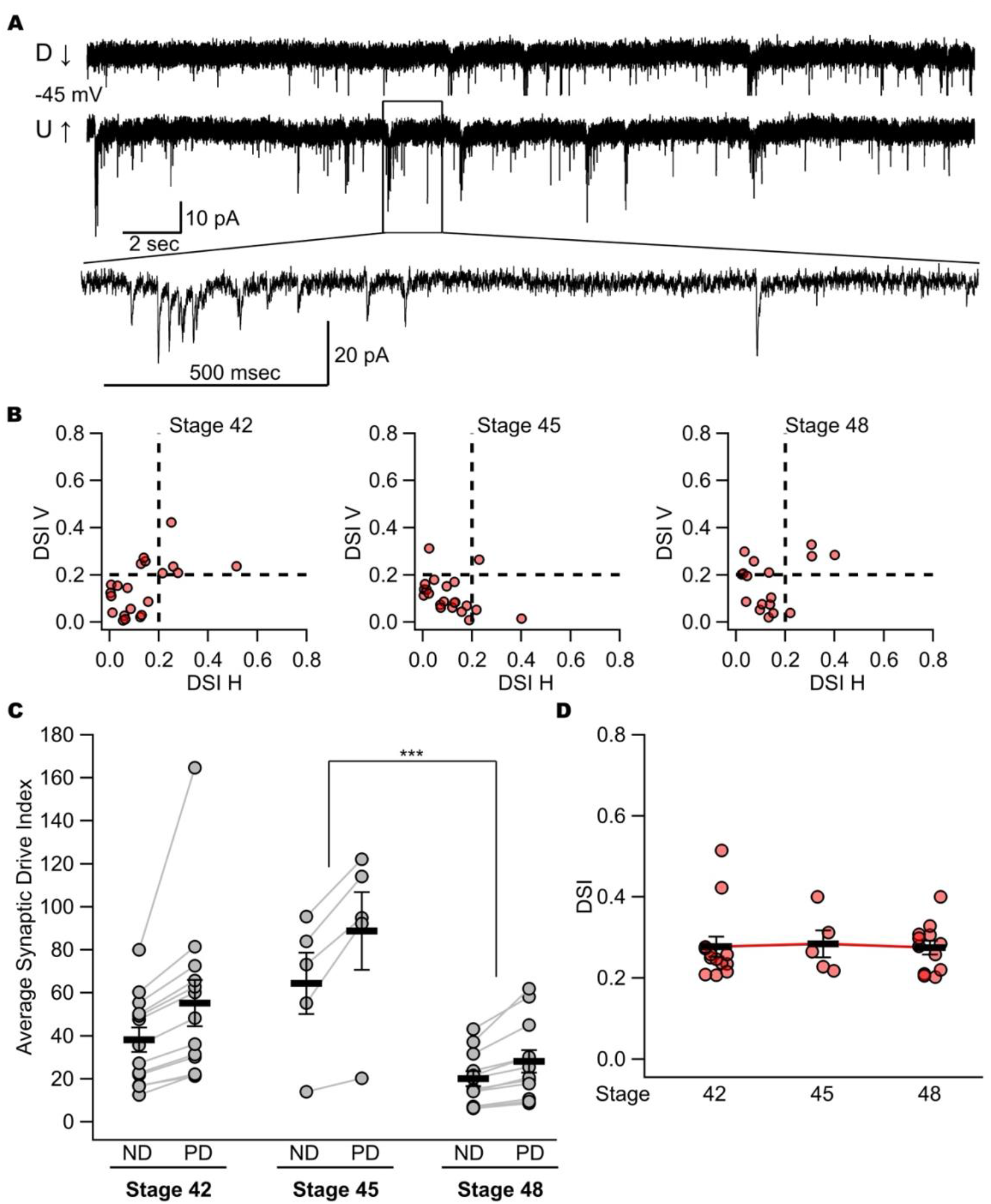
Tectal neurons do receive direction selective inputs, and the strength of them does not vary much across developmental stages. **(A)** 30 second chunk with a zoomed in 2 second snippet of a sample recording trace of a voltage-clamp recording recorded at -45mv, from a stage 42 direction selective tectal neuron, preferring the downward to upward direction. **(B)** Dot plots representing the relative strength of direction tuning, the DSI, of synaptic inputs that stage 42, 45, and 48 midbrain tectal neurons received when moving bars are projected to the contralateral eye. Each dot represents a single neuron that has met the events count criterion, displaying its DSI value for both vertical and horizontal orientations. The dash line represents the deliberate 0.2 DSI cut off. The proportion of DS tectal neurons fluctuated across development: stage 42: 38%, qualified cell n=21, 8 were DS; stage 45: 19%, qualified cell n=21, 4 were DS; stage 48: 53%, qualified cell n=17, 9 were DS. **(C)** Dot plot showing the average SDI of each tectal neuron, paired in non-preferred direction versus preferred direction across three developmental stages. For neurons displaying direction selectivity in both orientations, both pairs of their SDI values are included in the calculations. For between group comparisons, the SDI difference between stage 45 and stage 48 is statistically significant, \*\*\**p*<0.001, *p*=0.005, Mann-Whitney U test. The bars represent the SDI mean ± SEM. At stage 42, the average SDI for ND is 38.21±5.72; for PD is 55.27±10.78. At stage 45, the average SDI for ND is 64.38±14.24; for PD is 88.73±18.02. At stage 48, the average SDI for ND is 22.07±3.49; for PD is 28.08±5.23. **(D)** Categorical trend plot depicting the overall trend of input DSI for only direction selective tectal neurons across three key developmental stages. For neurons displaying direction selectivity in both orientations, both of their DSI values are included in the calculations. The average input DSI of tectal neurons at stage 42 is 0.28±0.02, n=13; at stage 45 is 0.28±0.03, n=5; and at stage 48 is 0.28±0.02, n=12. For between group comparisons, stage 42 vs. stage 45 tectal DSI is *p*=0.7028, n.s., Mann-Whitney U test; stage 45 vs. stage 48 tectal DSI is *p*=0.8048, n.s., Welch’s *t*-test. The bars represent the mean ± SEM. D↓: downward; U↑: upward; mV: millivolt; sec: seconds; msec: millisecond; pA: picoampere; DSI: direction selective index; V: vertical; H: horizontal; ND: non-preferred direction; PD: preferred direction.

### Direction selective output of tectal neurons sharpens across development

From a different set of tectal neurons, we measured direction selectivity of spiking output across the same three developmental stages. Figure 6A shows an example of a tectal neuron displaying DS output in the vertical orientation. While the percentage of tectal neurons that met the event count criterion was relatively low, most neurons that did meet the event count criterion displayed DS output (Table 1; Figure 6B). Figure 6C shows the average spike counts per 50 second trial elicited by the preferred and non-preferred directions of motion expressed by individual DS tectal neurons. Overall average spike frequencies exhibited a statistically significant decrease between developmental stages 45 and 48 (Figure 6C), while the average output DSI value exhibited a statistically significant increase (Figure 6D, Figure 4D-F). This increase in the average output DSI at developmental stage 48 was unexpected given that we observed no such increase in the average synaptic input DSI at this stage. This implies that developmental stage 48 tectal neurons transform their broadly direction tuned input to more narrowly tuned output.

**Figure 6:**
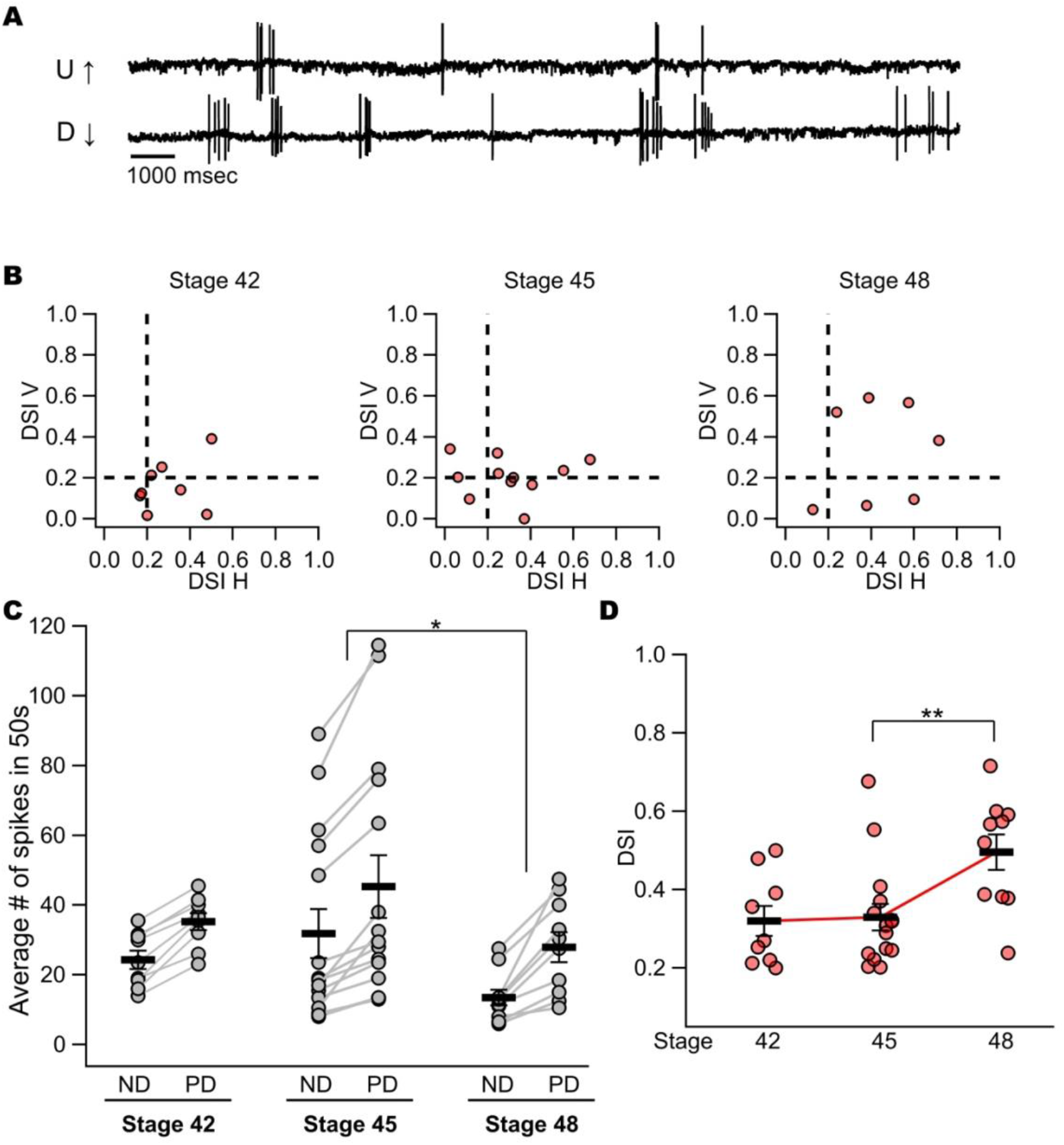
The output direction selectivity of tectal neurons exhibit a distinct developmental program across stages. **(A)** 20 second snippet sample trace of a loose cell-attached recording, from a stage 48 direction selective tectal neuron, preferring the upward to downward direction. **(B)** Dot plots representing the relative strength of direction tuning, the DSI, of output action potentials that stage 42, 45, and 48 midbrain tectal neurons fired when moving bars are projected to the contralateral eye. Each dot represents a single neuron that has met the event count criterion, displaying its DSI value for both vertical and horizontal orientations. The dash line represents the deliberate 0.2 DSI cut off. For neurons in the optic tectum, about 75% of tectal neurons are DS at stage 42, qualified n=8, 6 were DS; about 91% of tectal neurons are DS at stage 45, qualified n=11, 10 were DS; about 85% of tectal neurons are DS at stage 48, qualified n=7, 6 were DS. **(C)** Dot plot showing average number of spikes in 50 seconds of each tectal neuron, paired in non-preferred direction versus preferred direction across three developmental stages. For neurons displaying direction selectivity in both orientations, both pairs of their spike count per 50 seconds trials are included in the calculations. For between group comparisons, the spiking difference between stage 45 and stage 48 is statistically significant, \**p*<0.05, *p*=0.04, Mann-Whitney U test. The bars represent mean ± SEM of the average number of spikes in each 50 second trial. At stage 42, the average spiking for ND is 24.33±2.61; for PD is 35.28±2.41. At stage 45, the average spiking for ND is 31.80±7.08; for PD is 45.30±9.00. At stage 48, the average spiking for ND is 13.50±2.27; for PD is 27.95±4.26. **(D)** Categorical trend plot depicting the overall trend of output DSI for only direction selective tectal neurons across three key developmental stages. For neurons displaying direction selectivity in both orientations, both of their DSI values are included in the calculations. The average overall DSI of tectal neurons at stage 42 is 0.32±0.04, n=9; at stage 45 is 0.33±0.03, n=15; and at stage 48 is 0.50±0.05, n=10. For between group comparisons, stage 48 vs. stage 45 tectal DSI is \*\**p*<0.01, *p*=0.0054, Mann-Whitney U test; stage 42 vs. stage 45 tectal DSI is insignificant, *p*=0.60, Mann-Whitney U test. The bars represent the mean ± SEM. U↑: upward; D↓: downward; msec: millisecond; DSI: direction selective index; V: vertical; H: horizontal; s: seconds; ND: non-preferred direction; PD: preferred direction.

### Sharpening of tectal spike output at developmental stage 48 does not appear to be manifested via changes in synaptic input

Because tectal input tuning did not sharpen across development (Figure 2D), we asked whether the sharpening of tectal output at stage 48 could arise from developmental changes in activity-dependent refinement. The retinotectal circuit undergoes NMDA receptor-dependent refinement during development (Cline and Constantine-Paton,1990; O’Rourke et al.,1994; Rajan and Cline,1998; Dong et al., 2009; Hamodi et al., 2016). While we did not observe a sharpening of direction tuning of synaptic input received by tectal neurons across development, suggesting a lack of refinement of DS input, we still wanted to confirm that the sharpening of DS output was not manifested by any NMDA receptor-mediated synaptic refinement. To test this, tadpoles were exposed to the NMDAR blocker APV beginning at stage 45 (Dong et al., 2009; Udoh et al., 2025), and *in vivo* loose cell-attached recordings at stage 48 were carried out to measure output direction selectivity (Figure 7A). We found that chronic NMDA receptor blockade across stage 45 and 48 elicited a statistically insignificant effect on the average strength of DS output relative to batch-matched controls (Figure 7B). These data indicate that the developmental sharpening of DS spike output is not due to NMDAR dependent refinement of synaptic input.

**Figure 7:**
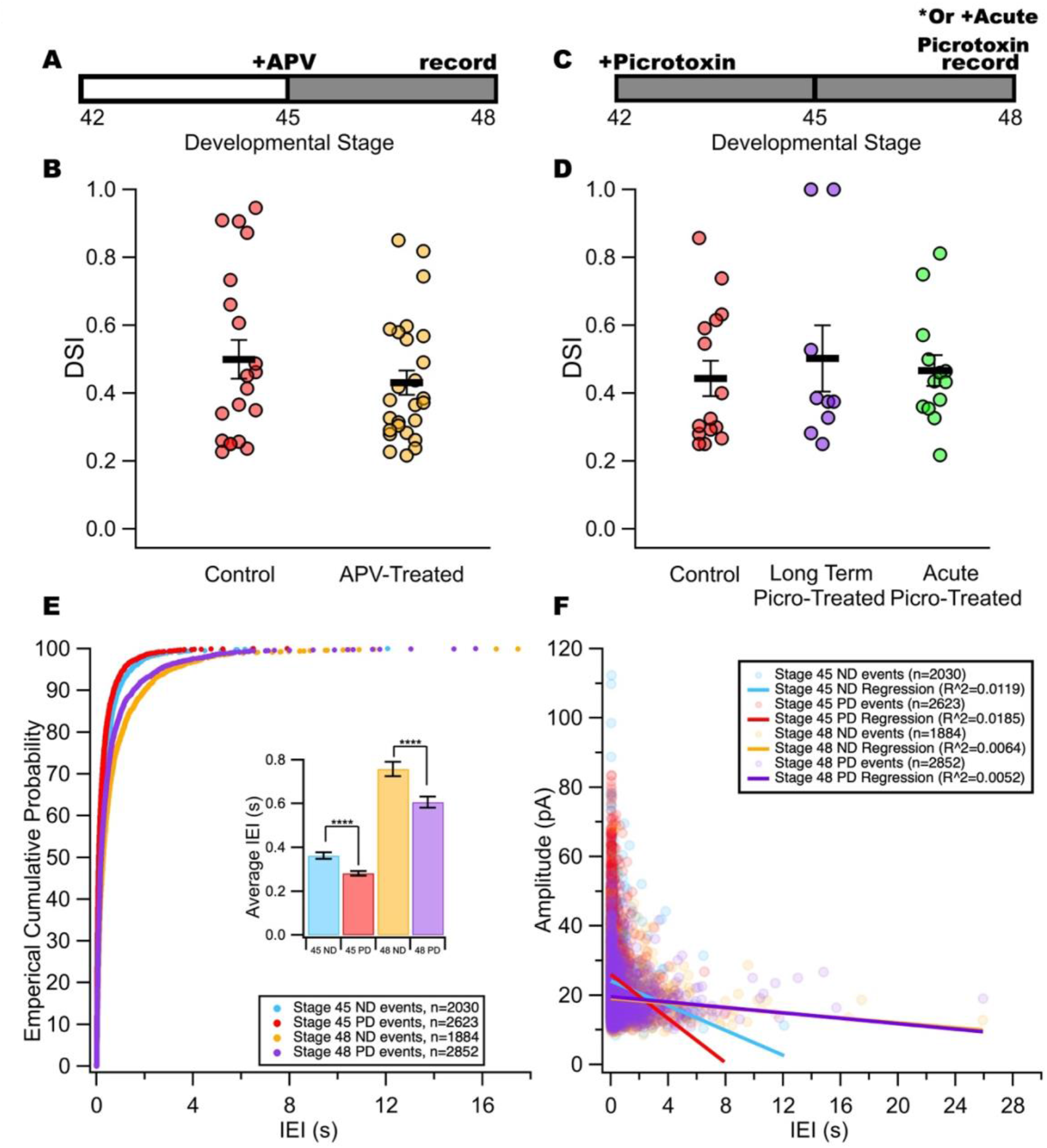
Sharpening of stage 48 tectal neurons’ direction selective output is neither due to NMDAR nor inhibition, but shorter IEI could contribute to it. **(A)** Schematic showing the timeline of the APV experiments carried out. 100μM of APV were infused in the 10% Steinberg’s rearing solution of a selected few tadpoles at developmental stage 45, and *in vivo* loose cell-attached electrophysiological recordings in the tectum were carried out at developmental stage 48 for both batch-matched controls and APV treated tadpoles. **(B)** Dot plot representing the relative strength of direction tuning, the DSI, calculated from output action potential spiking fired by stage 48 batch-matched control tectal neurons in red, and APV treated tectal neurons in orange. For neurons displaying direction selectivity in both orientations, both of their DSI values are included in the calculations. Batch-matched control stage 48 tectal neuron average DSI is 0.50±0.06, n=20; APV-treated stage 48 tectal neuron average DSI is 0.43±0.04, n=26; APV versus control, n.s., *p*=0.5349, Mann-Whitney U test. The bars represent the mean ± SEM. **(C)** Schematic showing the timeline of the picrotoxin experiments carried out. For long term picrotoxin treatment experiment, 100µM of picrotoxin dissolved in 100% DMSO were infused in the 10% Steinberg’s rearing solution of a selected few tadpoles at developmental stage 42, and *in vivo* loose cell-attached electrophysiological recordings in the tectum were carried out at developmental stage 48 for both batch-matched controls and long term picrotoxin treated tadpoles; For acute picrotoxin exposure experiment, 100µM of picrotoxin dissolved in 100% DMSO were infused in the external recording solution at stage 48 before recording in *in vivo* loose cell-attached configuration. Batch-matched controls for acute picrotoxin were also recorded from *in vivo* loose cell-attached configuration. **(D)** Dot plot representing the relative strength of direction tuning, the DSI, calculated from output action potential spiking fired by stage 48 batch-matched control tectal neurons in red, long term picrotoxin treated tectal neurons in purple, and acute picrotoxin treated tectal neurons in green. For neurons displaying direction selectivity in both orientations, both of their DSI values are included in the calculations. Since the result of batch-matched controls between long term and acute picrotoxin treatment had no statistical differences, two groups of controls were combined in the final analysis. Batch-matched control stage 48 tectal neuron average DSI is 0.44±0.05, n=15; long term picrotoxin treated stage 48 tectal neuron average DSI is 0.50±0.10, n=9; versus control: n.s., *p*=0.6288, Mann-Whitney U test; acute picrotoxin treated stage 48 tectal neuron average DSI is 0.47±0.05, n=13; versus control: n.s., *p*=0.4324, Mann-Whitney U test. The bars represent the mean ± SEM. **(E)** Empirical percentage cumulative distribution plot of the preceding IEIs for stage 45 and stage 48 direction selective neurons preferred and non-preferred direction inputs. The horizontal axis shows the interval preceding each event. Inset is a bar graph of the same data showing the overall average IEI in seconds of the respective groups (\*\*\*\**p*<0.0001, Mann-Whitney U test, Stage 45: non-preferred direction, n=2030; preferred direction, n=2623. Stage 48: non-preferred direction, n=1884; preferred direction, n=2852). The bars represent the mean ± SEM. Stage 45 non-preferred direction labeled in blue; stage 45 preferred direction labeled in red; stage 48 non-preferred direction labeled in orange; stage 48 preferred direction labeled in purple. **(F)** Dot plot showing relationship between inter-event-interval and amplitude of tectal neurons in stage 45 and stage 48 direction selective neurons preferred and non-preferred direction inputs. The correspondingly colored lines represent the curve fitted linear regression of each stage and direction’s IEI vs. amplitude correlations. APV: 2-amino-5-phosphonopentanoic acid; DSI: direction selective index; Picro: picrotoxin; IEI: inter-event-interval; s: seconds; ND: non-preferred direction; PD: preferred direction; pA: picoampere; R^2: coefficient of determination.

Because inhibitory synaptic input activated by the non-preferred direction of motion has been reported to tune direction selectivity in both the zebra fish optic tectum (Grama and Engert, 2012; Reviewed in Gebhardt et al., 2013), and the mammalian visual cortex (Li et al., 2015; Wilson et al., 2018), we next asked whether the maturation of inhibitory signaling may underlie the sharpening of DS output that we observed in stage 48 tectal neurons. To test this, the GABAa receptor blocker picrotoxin was added to the external recording solution and loose cell-attached recordings were carried out to measure spiking in response to the same four orthogonal directions of motion (Figure 7C). We found that DSI values obtained from recordings carried out in the presence of the GABA receptor blocker were statistically indistinguishable from batch-matched controls (Figure 7D), indicating that GABA-mediated inhibitory currents do not contribute to the observed sharpening of DS output at developmental stage 48. To determine whether the increase in output direction tuning may require long-term GABA-mediated synaptic transmission, picrotoxin was added to the rearing solution at developmental stage 42, and then loose cell attached recordings carried out at developmental stage 48 (Figure 7D). We observed that chronic GABA receptor blockade did not interfere with the sharpening of DS output at developmental stage 48 (Figure 7D).

Next, we asked whether changes in the *pattern* of synaptic input evoked by the preferred and non-preferred directions of motion may account for the significant increase in the average spike output DSI that we observe at developmental stage 48. The synaptic parameters that were measured to calculate the DSI (EPSC frequency and average amplitude) provide an estimate of the overall amount of synaptic drive elicited by the different directions of motion, but they do not reflect the temporal pattern of the input. For instance, two stimuli may evoke the same average frequency of events, but their temporal pattern will determine whether the events sum. To determine whether the pattern of synaptic input may differ between the preferred and non-preferred direction at developmental stages 45 and 48, we measured inter-event-intervals (IEIs) between synaptic events. This analysis shows that, at both these developmental stages, synaptic events elicited by the preferred direction of motion displayed shorter IEI’s compared to synaptic events evoked by the non-preferred direction (Figure 7E). Theoretically, this pattern of synaptic input generated by the preferred direction of motion would enhance the probability of summing of synaptic events, which would be more effective at depolarizing the neuron to reach spike threshold. Hence, in addition to the greater synaptic drive elicited by the preferred direction of motion, the shorter average IEIs of the synaptic events would also work to boost spike frequency. Because the different temporal patterns of synaptic input evoked by preferred and non-preferred directions of motion are common to both development stage 45 and 48, this cannot solely account for the significant increase in the average DSI value of spike output that we observe at stage 48. Next, we asked whether there may be a relationship between the IEI and EPSC amplitude. Plotting EPSC amplitude as a function of IEI reveals a weak negative correlation between these two parameters, indicating a tendency for higher-amplitude EPSCs to follow shorter IEIs, possibly due to facilitation (Figure 7F). This trend was observed to be more pronounced at stage 45 compared to stage 48 (Figure 7F) which further supports the notion that this pattern is due to facilitation given that the amount of facilitation expressed by retinotectal synapses is known to decrease over these same developmental stages (Aizenman and Cline, 2007). This suggests that the relatively weaker relationship between IEI and EPSC amplitude observed at developmental stage 48 cannot account for the increase in direction tuning of spike output.

All together, these results indicate that the observed increase in tectal neuron output direction selectivity between developmental stages 45 and 48 cannot be attributed to NMDA receptor-mediated refinement of DS input, changes in GABA receptor-mediated inhibitory signaling, nor a pattern of synaptic input evoked by the preferred versus non-preferred direction which is unique to developmental stage 48.

### Increase in spike threshold at developmental stage 48

How, then, is the broadly tuned DS synaptic input transformed into more sharply tuned output in the optic tectum? We reasoned that if the sharper direction tuning of developmental stage 48 tectal neuron output is not manifested through sharpening of direction selectivity of synaptic input, then it might instead involve a change in the tectal neuron’s intrinsic excitability – the ability of the neuron to fire action potentials. To investigate this, we analyzed voltage-dependent Na^+^ and K^+^ currents that had been collected from the tectal neurons before recording their synaptic responses to the moving bars of light (example Figure 8A). The current-voltage (IV) plots generated from these recordings reveal a decrease in average maximum peak sodium currents across development, at every voltage step that activates this current (Figure 8B). Averaging the tectal neurons’ maximum peak Na+ current (the maximum peak current, regardless of which voltage step at which it occurred) shows a statistically significant decrease in average maximum peak Na+ currents between developmental stage 45 and 48 (Figure 8C; \*\**p*=0.0013; n=21 for stage 45, n=19 for stage 48), a finding previously reported by Pratt and Aizenman (2007). No statistically significant differences in the average peak K+ current at the beginning of the voltage step which is comprised of the transient and persistent components (the peak current evoked by the last, most depolarizing step of the I-V protocol) were detected across development. To further test whether the observed decrease in the average maximum peak Na^+^ current in stage 48 tectal neurons was sufficient to decrease intrinsic excitability, we also compared action potential (spike) thresholds of developmental stage 45 and 48 tectal neurons. Indeed, spike thresholds exhibited a statistically significant increase in stage 48 tectal neurons (Figure 9; \*\*\**p*=0.0003; n=21 for stage 45, n=16 for stage 48), suggesting an overall decrease in intrinsic excitability. Two additional lines of evidence are consistent with a decrease in intrinsic excitability at developmental stage 48: (1) the percentage of tectal neurons that met the event count criterion reached an all-time low of 10% at developmental stage 48 (Table 1), (2) the developmental stage 48 neurons that did meet the event count criterion displayed an overall lower average spike frequencies associated with both the preferred and non-preferred directions of motion compared to those at developmental stage 45 (Figure 6C).

**Figure 8:**
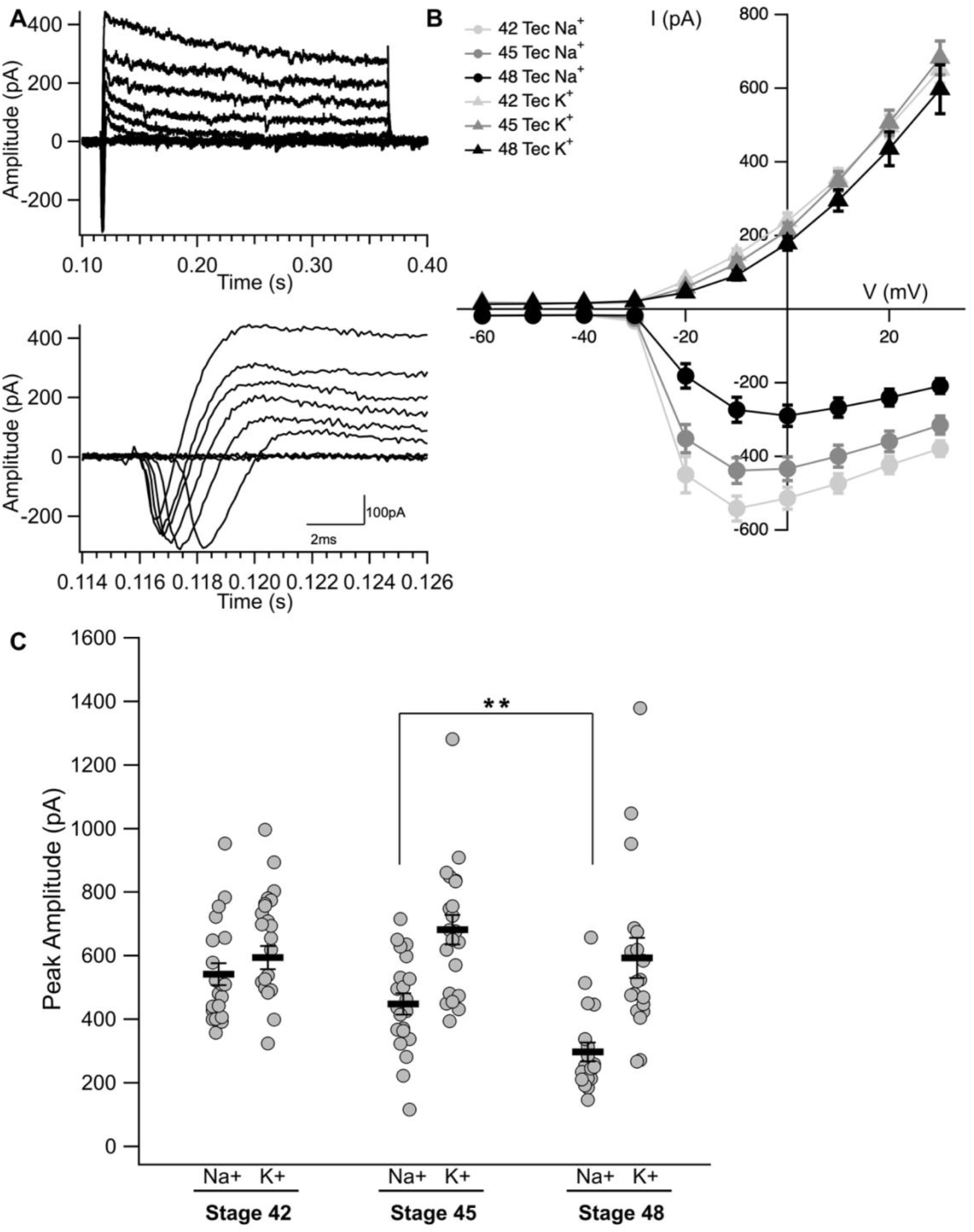
Tectal neurons exhibit a decreased intrinsic excitability across developmental stages. **(A)** Sample of a voltage-clamp recording of mixed currents collected from a stage 48 tectal neuron. The lower panel provides a zoomed in focus of the peak Na^+^ current of the same neuron. Traces represent currents activated in response to increasingly depolarizing steps. Currents were elicited by a series of depolarizing voltage steps from a holding potential of -60mV. The inward, inactivating portion of each trace represents Na^+^ current. The outward portion of each trace represents a transient IA-type and slower, persistent outward K^+^ currents. **(B)** I-V plot displaying average maximum peak voltage-gated Na^+^ (downward inflection) and average maximum peak K^+^ (upward) currents as a function of voltage steps for tectal neurons across three developmental stages. The maximum peak K+ currents reflect the fast, transient I_A_ type K+ current. **(C)** Categorical dot plot showing the average maximum peak amplitude of Na^+^ and K^+^ currents of tectal neurons across developmental stages. Na^+^ currents significantly decrease between stage 45 and stage 48: stage 45 average peak Na^+^=-448.2±33.04 pA; stage 48 average peak Na^+^=-297.6±30.01 pA; \*\**p*<0.01, *p*=0.0013, Mann-Whitney U test. The bars represent the mean ± SEM. I-V: current-voltage relation; pA: picoampere; s: second; I: currents; V: voltage; mV: millivolt; Na^+^: sodium currents; K^+^: potassium currents.

**Figure 9:**
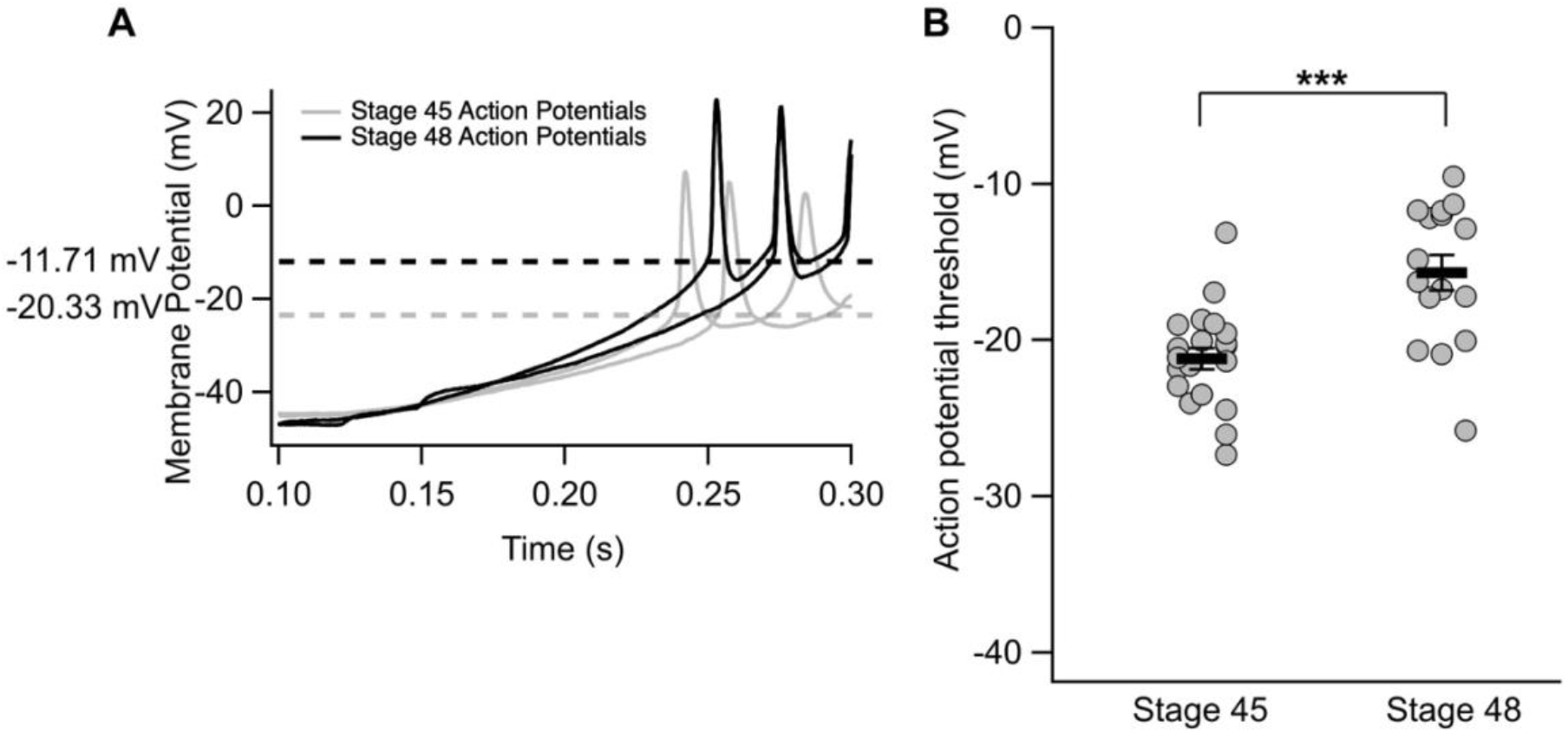
Stage 48 tectal neurons have higher action potential thresholds than stage 45 tectal neurons. **(A)** Samples of stage 45 and stage 48 tectal neurons’ AP threshold elicited by the current-ramp test protocol. **(B)** Dot plot showing the overall distribution of each recorded stage 45 and stage 48 tectal neurons AP threshold. There was a statistically significant difference between stage 45 tectal neurons AP threshold and stage 48 tectal neurons AP threshold: stage 45 mean=-21.19±0.69 mV; stage 48 mean=-15.68±1.12 mV; \*\*\**p*<0.001, *p*=0.0003, Welch’s *t*-test. The bars represent the mean ± SEM. AP: action potential; s: seconds; mV: millivolt; s: seconds.

We next asked whether the *magnitude* of the decrease in average spike frequencies between stage 45 and 48 may be greater for spiking evoked by the non-preferred direction of motion. Such an asymmetric effect could account for the observed sharpening of output direction tuning at the later stage. Consistent with this possibility, we observed that the average spike frequencies evoked by the preferred and non-preferred direction of motion decreased by 35% and 58%, respectively, indicating that the increase in spike threshold had a proportionally greater impact on spike output evoked by the non-preferred direction. Statistical analysis, however, indicated that the decrease in average spike frequency evoked by neither the preferred nor non-preferred direction was significant. This suggests that while the higher spike threshold contributes to increasing the difference between average spike frequencies evoked by the preferred and non-preferred direction and thus increasing the average spike DSI, it is probably not the only contributor.

In summary, we have identified three major changes that occur between developmental stage 45 and 48 in the optic tectum: (1) a steep decline in the overall strength of synaptic drive received by tectal neurons in response to the moving bar stimulus, (2) a significant increase in their spike threshold and concomitant decrease in spike frequencies in response to the moving bar stimulus, and (3) a sharpening of spike direction tuning in the absence of sharpening of synaptic input direction tuning.

## Discussion

Previously we compared the functional development of the retinotegmental and retinotectal projections across three key stages of visual system development (Udoh et al., 2025). Here, we extend this comparative approach, this time focusing on development of direction selectivity of neurons residing in these two retinorecipient visual processing centers of the midbrain. We found that in the basal optic nucleus of the midbrain tegmentum, direction selectivity of both synaptic input and spike output appears hard-wired and stable *ab initio*. In the optic tectum, the average degree of direction tuning (DSI values) remained stable for the synaptic input while tuning of spike output increased between developmental stages 45 and 48. This gave rise to the question of how direction tuning of spike output became more selective without an increase direction tuning of synaptic input. Our data suggest that a naturally occurring increase in spike threshold at developmental stage 48 filters spike output in a way that amplifies the difference between the spike frequencies evoked by the preferred and non-preferred direction of motion, thereby rendering the output more sharply tuned.

### Tegmental neurons display direction selectivity that is moderate in strength and stable across development of the retinotegmental projection

We measured direction selectivity of both the synaptic input received by tegmental neurons and the direction selectivity of their spiking (action potential) output from developmental stage 42, the time in development when the retinal ganglion cell axonal inputs have just started to form functional synapses with the postsynaptic tegmental neurons, forming the AOS. We found that neither the percentage of tegmental neurons that qualified as direction selective nor the strength of their direction selectivity appeared to change over development (Figures 2 and 3). This stable strength of direction selectivity across development is similar to our previous finding that the retinotegmental projection (the AOS) functions robustly as soon as the projection forms, and does not appear to strengthen or refine over development (Udoh et al., 2025). Furthermore, the average strength of direction tuning of synaptic input was as strong at developmental stage 42 as it was later in development, suggesting that tegmental neurons of the amphibian AOS inherit direction selectivity directly from direction selective retinal ganglion cells (RGC’s). This concept is consistent with previous studies in the mouse which report that specific classes of DS RGC’s directly synapse onto neurons of the AOS nuclei in the midbrain tegmentum (Yonehara et al., 2009; Sun et al, 2015). Finally, the moderate strength of direction tuning displayed by tegmental neurons means that while these neurons are most strongly activated by the preferred direction of motion, they also receive relatively strong input in response to the opposite direction. Relatively broad direction tuning has been described for tegmental AOS neurons in the adult frog, bird, and mammal (Fredericks et al., 1988; Crowder et al., 2003; Elstrott and Feller, 2009; Bastakov et al., 2015; Reviewed in Cang et al., 2018)

### Percentage of tectal neurons that receive direction selective synaptic input appears to increase dramatically between developmental stage 45 and 48

The development of direction selectivity in the optic tectum appeared to be more dynamic compared to that of the midbrain tegmentum. First, between developmental stages 42 and 45, we observed a marked decline in the percent of tectal neurons which received direction selective input, and then, between developmental stages 45 and 48, we observed a steep increase (Table 1). What may account for this transient decline in the percentage of neurons receiving DS input at developmental stage 45? We know that at developmental stage 45, intrinsic excitability is relatively high (spike thresholds are low). This would render the population of tectal neurons more likely to fire action potentials in response to both direct RGC input as well as RGC-evoked recurrent (tectal-tectal) input. At this stage, the recurrent activity has not yet refined, and compared to developmental stage 48, is known to be especially long-lasting (Pratt et al., 2008). Hence, motion evoked RGC input to the highly excitable optic tectum could theoretically generate a sufficiently large and long-lasting volume of synaptic activity, effectively masking any differences in the synaptic events generated by the preferred versus non preferred direction of motion. Our data support this scenario: at stage 45, the percentage of neurons whose input qualifies as direction selective is at its trough, while the average synaptic drive index is at its peak (Figure 5B, 5C; Table 1). The decrease in intrinsic excitability and rise in spike threshold that we observe at developmental stage 48 appears to dampen overall RGC-evoked activity in the optic tectum. Indeed, both the overall spike frequencies and amount of synaptic drive received by tectal neurons significantly decreased at this stage. This implies that only relatively strong, synchronized synaptic input, input that is driven by the moving stimulus, will be capable of inducing spiking. Consistent with the notion, it has been shown in the mouse superior colliculus that direction tuning is sharpened by the refinement of local recurrent circuitry (Shi et al., 2017).

Another possibility is that the steep increase in the percentage of neurons that receive DS input between stage 45 and 48 could be due to synaptic refinement. It is well established that the retinotopic map which forms across the optic tectum refines across these same two stages of development. The refinement of this map leads to more spatially compressed receptive fields, and the consequence is improved visual acuity (Dong et al., 2009; Hiramoto and Cline, 2014). Assuming that direction selectivity in the tectum is inherited from DS retinal ganglion cells, then perhaps a majority of tectal neurons which are destined to be direction selective may receive input from a broad range of DS RGCs early on and then refined such that any given tectal neuron receives input predominantly from DS RGCs that are selective for the same direction. This theory is supported by a previous study on the tadpole optic tectum which showed that developmental stage 42 and 45 tectal neurons can be entrained to be direction selective to any direction of motion, interpreted to mean that tectal neurons are not predetermined to be direction-selective for any specific direction of motion (Engert et al., 2002). However, like refinement of the retinotopic map, this entrainment was found to involved an NMDA receptor-dependent mechanism, while our data suggest that the natural development of direction selectivity of tectal neurons does not require activation of NMDA receptors (Figure 7A, 7B). Hence, if the increase in the percentage of DS tectal neurons between developmental stages 45 and 48 is due to refinement of DS input, it is happening via a non-canonical NMDAR-independent mechanism.

While these mechanisms may account for the steep increase in percentage of tectal neurons that receive DS input, it is difficult to reconcile how they could increase the percentage DS tectal neurons without also increasing the strength of direction tuning (DSI values). Perhaps the limit on direction tuning of synaptic input is set by the direction tuning of the RGC input itself.

### The significant increase in spike threshold at developmental stage 48 is likely the intracellular mechanism which transforms relatively weak direction selectivity of synaptic input into more strongly selective output

We also observed, between developmental stages 45 and 48, a significant increase in the average strength of DS spike output in the absence of an increase in the average strength of DS synaptic input. In other visual systems, it has been reported that direction tuning is sharpened by inhibitory input activated by the non-preferred direction (Grama and Engert, 2012; Reviewed in Gebhardt et al., 2013; Li et al., 2015; Wilson et al., 2018). This mechanism would fit our data too, because it would sharpen direction tuning of spike output without affecting direction tuning of excitatory synaptic input. However, we found that blocking GABA-mediated synaptic currents had no effect on DSI values of spike output at developmental stage 48, suggesting that, at least at this time in development, local inhibitory synaptic input is not a contributor to the sharper direction tuning of spike output.

Instead, our data indicate that the naturally occurring decrease in tectal neuron intrinsic excitability which occurs between developmental stages 45 and 48 at least contributes to the observed increase in the strength of spike output direction selectivity. It has been long known that the spike threshold increases strength of direction selectivity by adding a nonlinear filter on the incoming DS input (Jagadeesh et al, 1997; Priebe and Ferster, 2008; Roy et al., 2020). We observe this phenomenon in our data: across all three developmental stages, the average strength of direction selectivity of both tegmental and tectal neuron spike output is consistently stronger (more tuned) than that of their corresponding synaptic input (Figures 2-3; Figures 5-6; Table 1). Between stage 45 and 48, however, this nonlinear filter becomes stronger due to a naturally occurring increase in spike threshold. Consistent with a higher spike threshold, we also observed an overall decrease in average spike frequencies in response to both the preferred and non-preferred directions of motion. Our data show that the average decrease in absolute number of spikes evoked by the preferred and non-preferred directions of motion were essentially the same (i.e. the average decreased by 17 spikes/ 50 second trial between stage 45 and 48 for the preferred direction; and 18 spikes/ 50 second trial for the non-preferred direction). But, subtracting essentially the same number of spikes from both the preferred and non-preferred direction of motion affects them differently: because the average number of spikes / 50 second trial evoked by the non-preferred direction is, by definition, significantly less than that evoked by the preferred direction, subtracting the same number of spikes from both directions has a greater impact on the spike frequency evoked by the non-preferred direction. In other words, the equal decrease in average absolute number of spikes generates unequal proportional effects (Figure 10).

**Figure 10:**
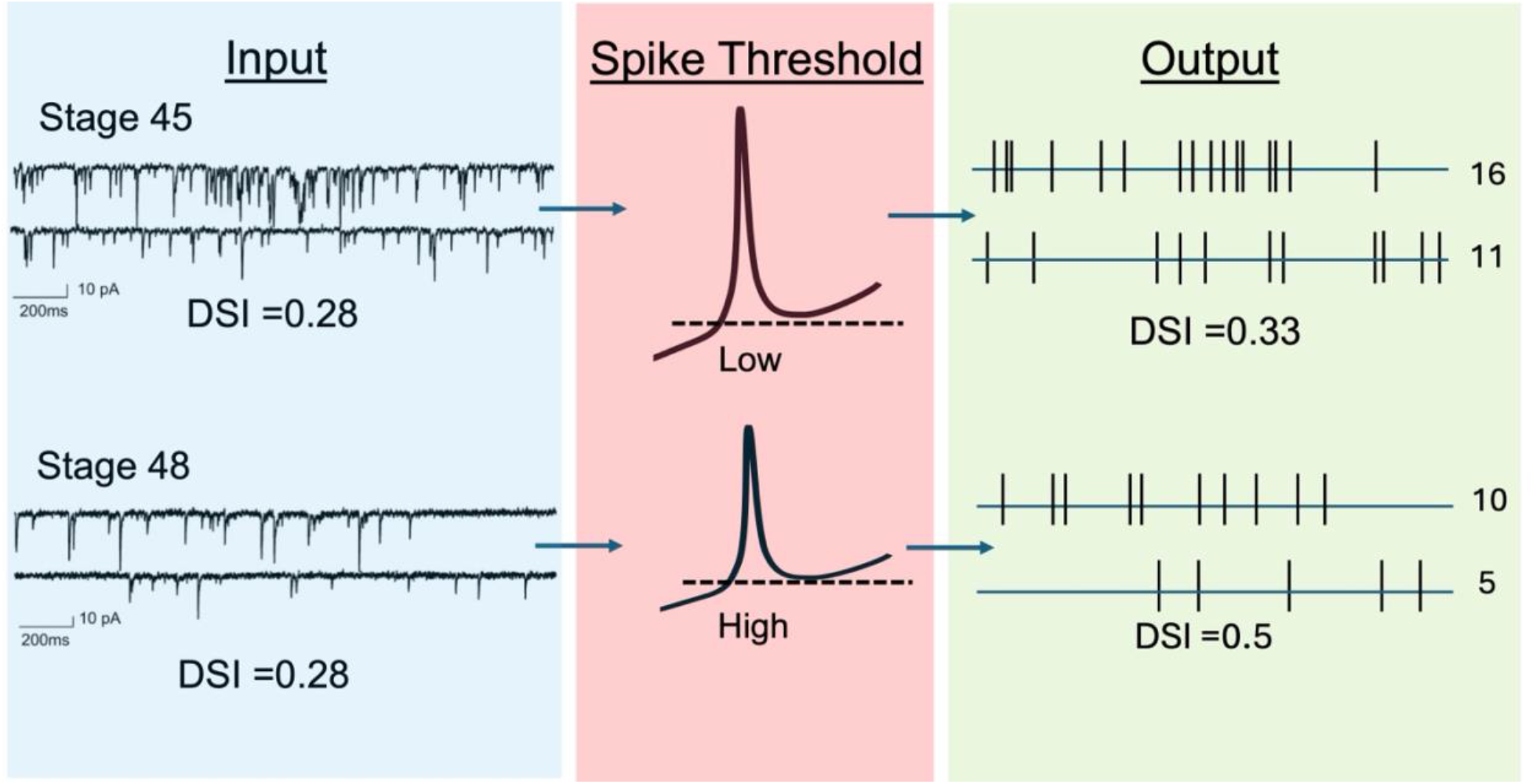
Theoretical model of the development of direction selectivity in the optic tectum. (Left) While overall amount of motion-evoked synaptic drive received by tectal neurons decreases between developmental stage 45 and 48, their DSI values are strikingly similar. (Middle) Spike threshold changes over development: relatively low at stage 45, high at stage 48. (Right) The relatively high spike threshold at developmental stage 48 decreases overall spike frequencies in both the preferred and non-preferred directions in a way that impacts the spike frequency evoked by the non-preferred direction to a greater degree, thus increasing direction tuning of the spike output.

The idea that changes in intrinsic excitability can tune direction selective spike output has also been described in a ferret study which compared direction selectivity in the visual cortex of naïve and experienced ferrets. The neurons of the experienced ferrets displayed stronger direction tuning due to an *increase* in intrinsic excitability (Roy et al., 2020). In this case, the increase in intrinsic excitability preferentially increased spiking in response to the preferred direction, which in turn increased DSI. Hence, in our study, we determined that a decrease in intrinsic excitability increases direction tuning by preferentially decreasing spiking rates in the non-preferred direction, and in the ferret study, it was determined that an increase in intrinsic excitability increases tuning by preferentially increasing spike rates in the preferred direction. This shows how the strength of direction selectivity can be achieved by shifting the nonlinear filter up or down to a degree that preferentially affects either the spiking in preferred or non-preferred direction of motion. This gives rise to interesting implications about the magnitude of changes in intrinsic excitability combined with specific strengths and patterns of synaptic input.

### A model and implications on relationship between strength and pattern of synaptic input and levels of intrinsic excitability

In summary, we have identified major changes in both levels of synaptic drive and intrinsic properties that occur between developmental stage 45 and 48 in the optic tectum: (1) a steep decline in the overall strength of synaptic drive received by tectal neurons in response to the moving bar stimulus, and (2) a significant increase in their spike threshold and concomitant decrease in spike frequencies. May the two naturally occurring changes in synaptic input and intrinsic excitability, combined, account for the significant increase in the average spike output DSI that is observed at developmental stage 48? Our results generated by a model neuron suggest they do. Notably, this model shows that neither the combination of stage 45 synaptic input with developmental stage 48 intrinsic excitability, nor stage 48 synaptic input with stage 45 intrinsic excitability generates a direction selectivity that is as strong as the combination of stage 48 synaptic input with stage 48 intrinsic excitability (Mondal et al., 2026).

The finding that a global decrease in intrinsic excitability can differentially affect spike frequencies elicited by the preferred and non-preferred directions gives rise to implications about the relationship between specific amounts of synaptic drive received by a given neuron and its level of intrinsic excitability. For instance, if all synaptic input were significantly stronger it is likely that the same increase in threshold that naturally occurs between developmental stages 45 and 48 would not impact firing rates in a way that would generate a higher DSI. And similarly, if the magnitude of the increase in spike threshold was beyond or below the naturally occurring increase, its effect on spike frequency also may not generate a greater DSI. This suggests that the natural activity levels of this biological circuit are balanced such that relatively small changes in input or intrinsic excitability levels can regulate how the circuit functions.

## Data Availability

The data that support the findings of this study are available upon request.

## Acknowledgements

The authors would like to thank Hollis Cline for her helpful comments and inspirations on the experimental design. This work was supported by an award from the National Science Foundation (Grant #2212591) to K.G.P., and an Institutional Development Award (IDeA) from the National Institute of General Medical Sciences of the National Institutes of Health (Grant #P20GM103432) to K.Z. and U.G.U., and the support of NIMH (R01MH046742) to Y.M. The authors would additionally like to recognize support from an Institutional Development Award (IDeA) from the National Institute of General Medical Sciences of the National Institutes of Health under grant number P20GM121310.

